# Weighted integration of short term memory and sensory signals in the oculomotor system

**DOI:** 10.1101/151381

**Authors:** Nicolas Deravet, Gunnar Blohm, Jean-Jacques Orban de Xivry, Philippe Lefèvre

**Affiliations:** Institute of Information and Communication Technologies, Electronics, and Applied Mathematics and Institute of Neuroscience, Université catholique de Louvain, B-1348 Louvain-La-Neuve, Belgium.; Centre for Neuroscience Studies, Queen’s University, Kingston, ON, Canada.; Canadian Action and Perception Network (CAPnet).; Department of Kinesiology, Movement Control and Neuroplasticity Research Group, Katholieke Universiteit Leuven, Leuven, Belgium

## Abstract

Oculomotor behaviors integrate sensory and prior information to overcome sensory-motor delays and noise. After much debate about this process, reliability-based integration has recently been proposed and several models of smooth pursuit now include recurrent Bayesian integration or Kalman filtering. However, there is a lack of behavioral evidence supporting these theoretical predictions.

Here, we independently manipulated the reliability of visual and prior information in a smooth pursuit task. Our results show that both smooth pursuit eye velocity and catch-up saccade amplitude were modulated by visual and prior information reliability.

We interpret these findings as the continuous reliability-based integration of a short-term memory of target motion with visual information, which support modelling work. Furthermore, we suggest that saccadic and pursuit systems share this short-term memory. We propose that this short-term memory of target motion is quickly built and continuously updated, and constitutes a general building-block present in all sensorimotor systems.

## Introduction

Over the last two decades, Bayesian integration of different signals (i.e. the weighted summation based on the respective reliability of each signal) has been widely applied to the study of cognitive processes. Its intrinsic ability to handle the uncertainty of different signals and combine them makes it a particularly useful tool to model how the brain handles the imperfect sensory representations of the external world. Indeed, be it relative to movement (Tassinari, Hudson, & Landy, 2006; Yang, Lee, & Lisberger, 2012), learning (Nassar, Wilson, Heasly, & Gold, 2010) or estimation (Stocker & Simoncelli, 2006), numerous studies have highlighted behaviors that exhibit reliability-based integration. Furthermore, when several senses give information about the same event or object, reliability based integration allows their (near) optimal combination (Ernst & Banks, 2002; Jacobs & Fine, 1999; Landy, Banks, & Knill, 2011). It has recently been suggested that internal predictive and sensory afferent information is combined in a Bayes-optimal fashion during smooth pursuit eye movements; however this prediction has not been tested explicitly.

The oculomotor system is well-studied and offers many typical examples of sensorimotor processes relying on both noisy sensory inputs (Bogadhi, Montagnini, & Masson, 2013; Osborne, Lisberger, & Bialek, 2005; Spering, Kerzel, Braun, Hawken, & Gegenfurtner, 2005) and prior experience (Kowler, Martins, & Pavel, 1984; Madelain & Krauzlis, 2003; Yang et al., 2012) to produce accurate movements. For example, visual tracking has to cope with sensory delays (Osborne, Bialek, & Lisberger, 2004) and internal noise (Osborne et al., 2005). By integrating visual inputs with past experience and cues, the oculomotor system can overcome sensory delays and noise to produce eye movements matching current target movement. In the pursuit system, this allows, for example, anticipatory smooth eye movements (Dodge, Travis, & Fox, 1930; Hayhoe, McKinney, Chajka, & Pelz, 2012; Kowler, Aitkin, Ross, Santos, & Zhao, 2014; Westheimer, 1954) or zero-lag smooth pursuit tracking of a sinusoidal target motion (Dodge et al., 1930; Orban de Xivry, Coppe, Blohm, & Lefèvre, 2013). In the saccadic system, predictive saccades can be observed when tracking a target jumping at a fixed frequency (Shelhamer & Joiner, 2003), or a bouncing ball (Diaz, Cooper, Rothkopf, & Hayhoe, 2013). Thus predictive and sensory information interact to drive oculomotor behavior.

The process of integration of sensory and predictive signals is still up to debate. Several models have been proposed to explain how sensory inputs and predictive signals might be integrated during smooth pursuit: some have used a switching mechanism between predictive and sensory-feedback processes (Barnes, 2008; Bennett & Barnes, 2004), but in recent years several authors turned to reliability-based integration (Bogadhi, Montagnini, Mamassian, Perrinet, & Masson, 2011; Dimova & Denham, 2009; Freeman, Champion, & Warren, 2010; Montagnini, Mamassian, Perrinet, Castet, & Masson, 2007). In 2013, Orban de Xivry, Coppe et al. proposed a model that, for the first time, was able to simulate the integration of a continuous flow of sensory information with past experience to drive motor behavior. This model used reliability-based integration (Kalman filtering) both to predict target movement and to simultaneously build a dynamic memory of it. Specifically, the model predicts that the estimated target movement is the reliability-weighted (reliability = 1/variance) average of visual and predictive target motion signals, in agreement with Bayes-optimal cue integration. This model managed to reproduce a large repertoire of pursuit behaviors. The same year, Bogadhi, Montagnini and Masson (2013) proposed a similar model that used a static memory of target velocity and two recurrent Bayesian networks for sensory and predictive signal integration. However, while there are several studies on the influence of prior information on oculomotor tracking (Kowler, 2011; Kowler et al., 2014; Wende, Theunissen, & Missal, 2013; Yang et al., 2012), few have investigated the mechanisms of integration of such prior information with sensory information, and how the reliability of the memory and the sensory inputs can influence this integration.

Here, we present two target-eye-tracking experiments in which we independently manipulated the uncertainties of visual information and short-term memory. As predicted by the model of Orban de Xivry, Coppe et al. (2013), we observe reliability-based integration of a short-term memory of target motion with visual information during movement. Furthermore, we show that this integration occurs in two types of eye movements: smooth pursuit and catch-up saccades. Given the similarities of smooth pursuit with other cortical sensorimotor systems (Lisberger, 2015; Lynch & Tian, 2006) and recent studies of memory updating (Gershman, Radulescu, Norman, & Niv, 2014; Nassar et al., 2010), we believe that our results validate an important prediction; continuous reliability-based integration of current sensory information with working memory signals is a general principle that is likely to be part of all sensorimotor processes.

## Results

### Prior information about the target velocity biases smooth pursuit eye velocity

In this study, we varied the reliability of sensory and prior information in order to see how each of these information channels can influence the oculomotor response. First, we measured the effect of prior information on smooth pursuit by comparing eye velocity for a selection of catch and control trials (Fig. 8B). The selected trials had identical target velocities (for example 15°/s) during the current trial but different target velocities during the previous ones (for example 15°/s for control and 20°/s for catch trials). In addition, these trials were matched by trial number. Therefore, any difference in behavior between these catch and control trials has to be attributed to the influence of prior information on the oculomotor response. This comparison is illustrated on Figure 1A, which shows average eye velocity profiles (noisy target) for control and catch trials for different trial numbers. On this figure, a clear and long-lasting bias (up to 500ms after the target motion onset) can be seen. Furthermore, this bias appears to increase with the number of previous trials.

**Figure 1:**
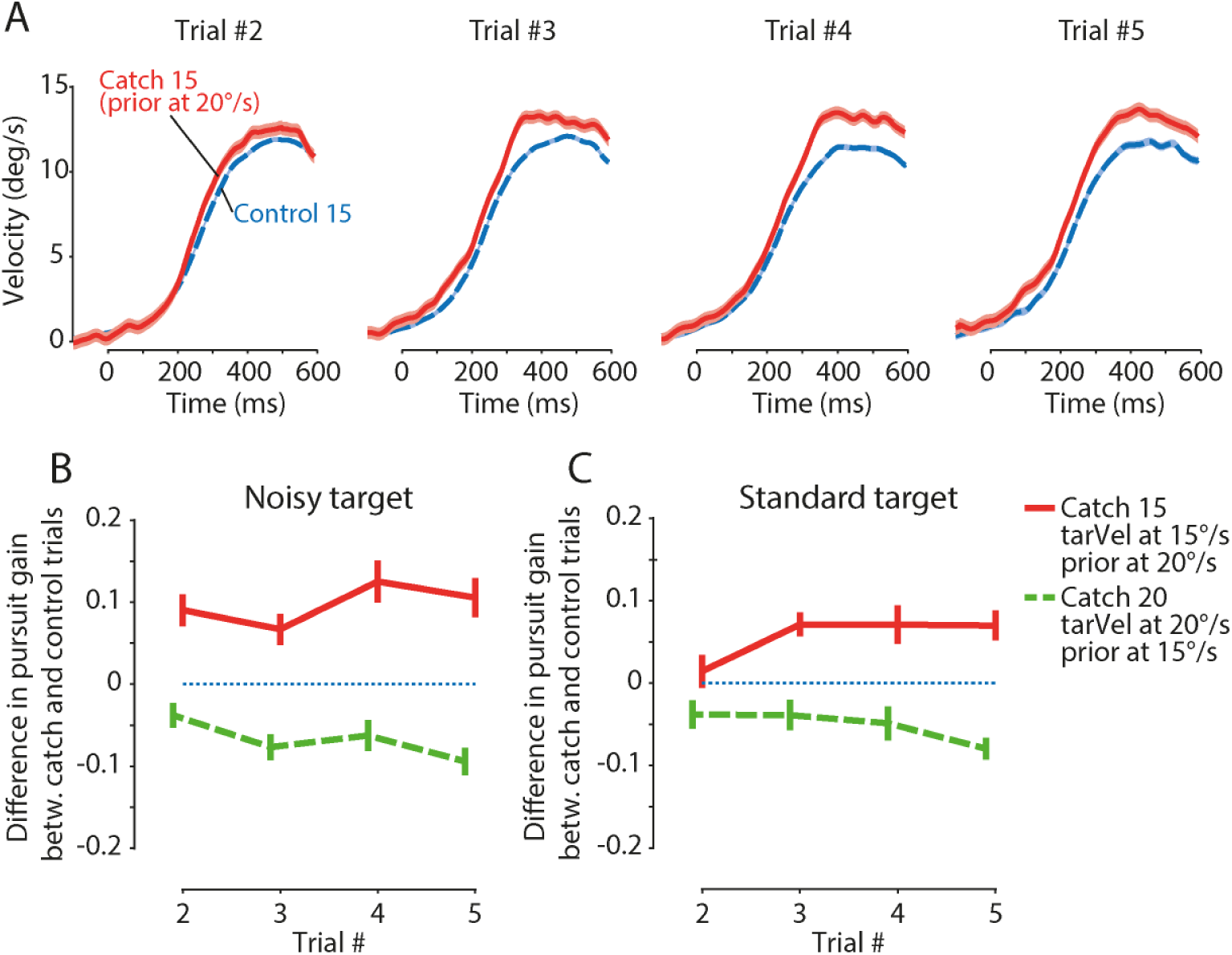
Prior information effect on eye velocity gain. **A.** Averaged eye velocity traces of Control15 and Catch15, for all participants (of the 1^st^ experiment, noisy target), for all trials allowing direct comparison between a catch and a control condition with a current target at 15°/s. Blue traces are the average eye velocity of control trials, while red traces are those of the catch trials. The surrounding hue indicates the standard error of the mean. **B.** Averages of participant’s mean differential gains of the smooth pursuit eye velocity response to a noisy target moving at 15°/s (full red line) or 20°/s (dashed green line) that was preceded by trials with the same target *either* at a *higher* (20°/s, full red line) velocity or a *lower* (15°/s, dashed green) velocity (catch trials). The error bars indicate the standard error of the mean. **C.** Same as panel B, but for the second experiment with a standard target. Dashed lines indicate a 15°/s prior target velocity.

To evaluate the influence of the prior information on the smooth pursuit response, we computed the steady-state smooth pursuit gain for each trial (see Methods) and subtracted the gain of the control trial from the gain of the corresponding catch trial (Fig. 1B and Fig. 1C). For all types of catch trials, the smooth pursuit response was biased towards the velocity of the preceding trials (main effect of trial type; Noisy Target, catch15: F(1,12)=51.81, p<0.0001, ges=0.17, catch20: F(1,12)=53.11, p<0.0001, ges=0.1; Standard Target, catch15: F(1,12)=21.38, p=0.0006, ges=0.07, catch20: F(1,12)=24.95, p=0.0031, ges=0.06). That is, on a given trial, the steady state smooth pursuit gain was higher (resp. lower) if this trial was preceded by trials with higher (resp. lower) target velocity. This demonstrates the effect of prior information on smooth pursuit eye velocity. This effect was true for both target types.

### Visual information about the target velocity affects smooth pursuit eye velocity

The impact of visual information on the smooth pursuit eye movements was measured by comparing catch and control trials that had the same prior information (same number of previous trials with the same target velocity), but different target velocities (see Fig. 8B).

Looking at eye velocity, this comparison (see Fig. 2A) showed that visual information had a clear effect: conditions having a higher target velocity showed, when compared to controls, a bias towards higher eye velocities and conditions having a lower target velocity showed a bias towards lower eye velocities. This effect was already present during the first active trial (#2), persisted during later trials, and could be observed as late as 500ms after the target onset.

**Figure 2:**
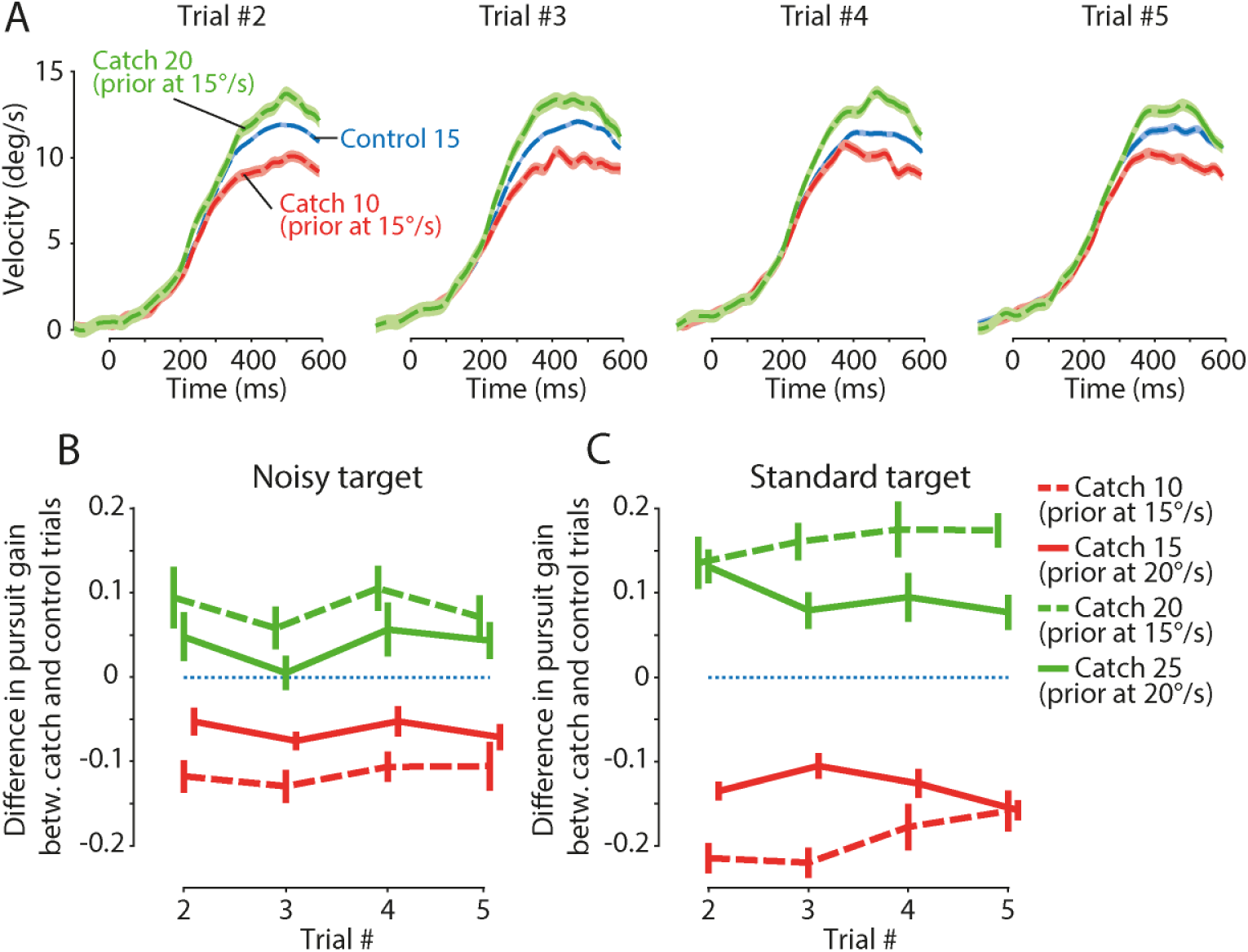
Visual information effect on eye velocity gain. **A.** Averaged velocity traces of all participants (noisy target), for all trials with the same target velocity prior – 15°/s – and all comparable trial numbers (all but the first and the last). The conditions Control15 (blue trace), Catch10 (decrease of velocity, red trace) and Catch20 (increase in velocity, green trace) have the same prior at 15°/s. The hue around each trace is the standard error of the mean. **B.** Averages of participant’s mean differential gains of the smooth pursuit eye velocity response to trials of a noisy target moving at 10°/s or 20°/s (resp. red and green dashed lines) *or* at 15°/s or 25°/s (resp. red and green solid lines) that were preceded by trials with the same target *either* at a *higher* velocity (all red lines) or a *lower* velocity (all green lines). The X-axis indicates the trial number and the error bars the standard error of the mean. **C.** Same as panel B, but for the second experiment with a standard target.

To highlight the effect of visual information on smooth pursuit eye velocity, we computed the difference of pursuit gains (see methods) to obtain the change in pursuit gain between catch and control trials. The differential gains show that the visual information modulated the smooth pursuit gain for all conditions of each experiment (Fig. 2B and C), which was confirmed by statistical analyses.

We found that the steady state eye velocity gains of catch trials were significantly different from those of control trials, both for the noisy target (main effect of trial type: F(2,24)=37.03, p<0.0001 Huynh-Feldt corrected, ges=0.23) and the standard target (main effect of trial type: F(2,24)=124.29, p<0.0001 Huynh-Feldt corrected, ges=0.53). Note that the interaction with the trial number is discussed in the last section of the results.

For both types of target, the magnitudes of the biases were higher for the conditions for which prior target velocity was at 15°/s (interaction effect between trial type and prior target velocity; Noisy target: F(2,24)=29.34, p<0.0001 Huynh-Feldt corrected, ges=0.03; Standard target: F(2,24)=30.08, p<0.0001 Huynh-Feldt corrected, ges=0.05). Thus overall, visual information affected pursuit gain.

### Saccade amplitude is biased by prior information about the target velocity

It is known that saccade and pursuit share many inputs. In particular, catch-up saccade amplitude is dependent on the difference between target and eye velocity (S. de Brouwer, Missal, Barnes, & Lefèvre, 2002). Therefore, if the internal representation of target velocity is biased by prior information, we would expect that this bias also results in alterations of catch-up saccade amplitude. Thus, we tested if the prior information about the target motion would also influence the catch-up saccades. To do so, we compared the amplitude of saccades in catch and control trials by computing the difference between the amplitude of saccades made during catch trials and the amplitude of saccades made during control trials of the same target velocity (see methods and Fig. 10).

We found that saccades were biased by prior information about target velocity; the amplitude of saccades made during catch trials was more likely to be larger than during control trials if the previous trials had a higher target velocity and to be smaller if the previous trial’s target was slower (main effect of trial type; Noisy Target, catch15: F(1,12)=34.47, p<0.0001, ges=0.28, catch20: F(1,12)=18.62 p=0.001, ges=0.18). When the standard target was presented on the screen, we found a similar effect of the prior on saccade amplitude (Standard target, catch15: F(1,12)=6.82, p=0.023, ges=0.02, catch20: F(1,12)=17.1, p=0.0014, ges=0.16). Fig. 3A illustrates this effect on some representative trials from one participant. Another way to look at this effect is presented in Fig. 10, where it can be observed that the data points from catch trials – the red dots – tend to be located above the regression line, indicating a bias towards larger saccades coherent with an effect of a higher target velocity prior.

**Figure 3:**
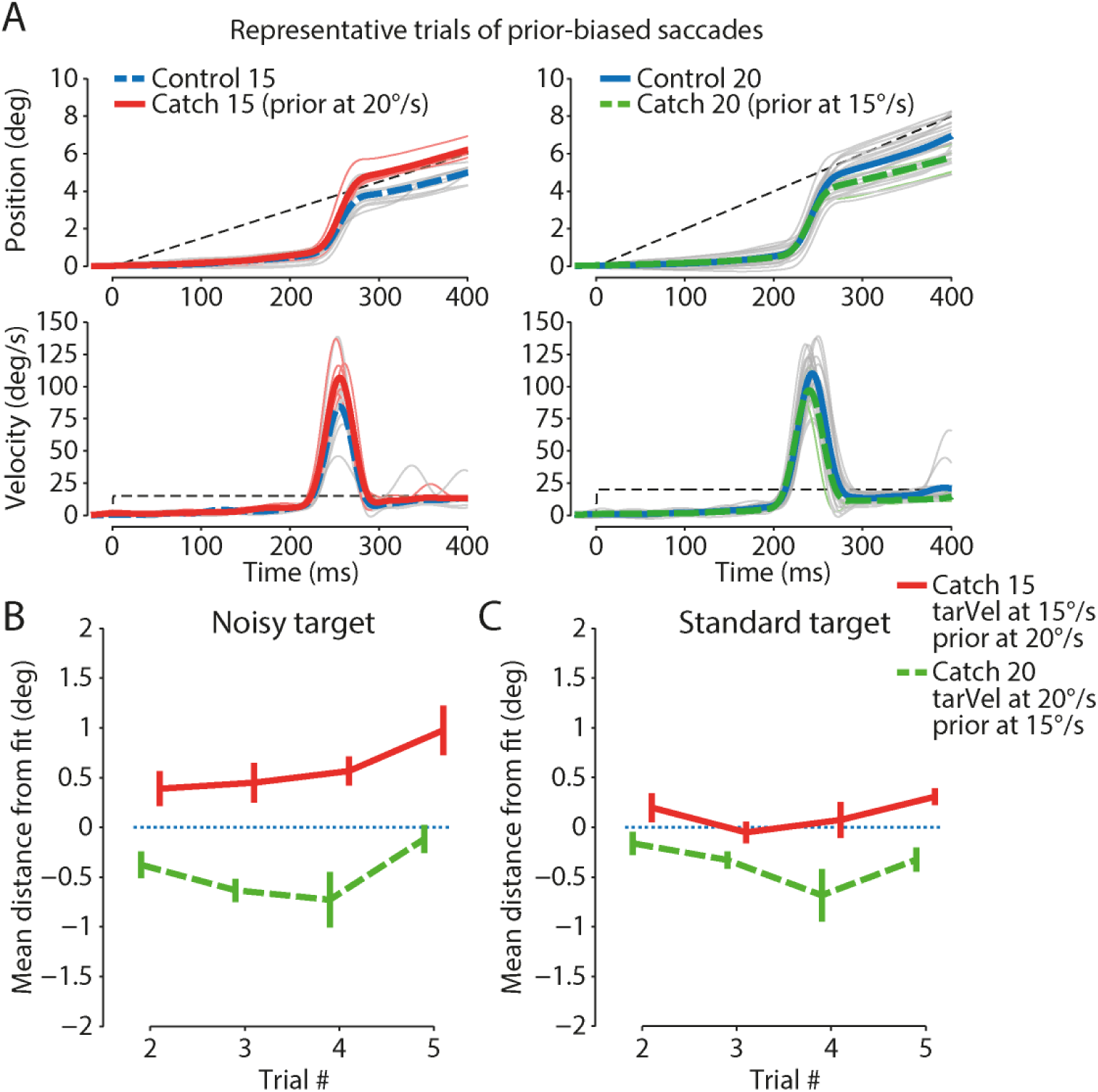
A. Effect of prior information on concomitant saccades. Position and velocity profiles of saccades made at the same time (15ms window) in representative control and catch trials. Left panel: saccades made around 220ms after the onset of the target, during control trials with a prior of target velocity at 15°/s (blue traces) and catch trials with a prior of target velocity at 20°/s (red traces). Right panel: saccades made around 220ms after the onset of the target, during control trials with a prior of 20°/s (blue traces) and catch trials with a prior of 15°/s (green traces). **B. Effect of prior information on saccades (noisy target).** Averages of participant’s mean residuals (vertical distance from fit on control data) of the amplitude of saccades made during catch trials, with respect to the trial number. The blue dotted line indicates the control reference (that has been subtracted from all data), red/green traces show residuals of saccades made during catch trials. The error bars indicate the standard error of the mean. **C.** Same as panel B, but for the second experiment with a standard target.

Because saccades can sometimes be adjusted online via internal feedback, we also studied the peak velocity of the saccades, as it is a good marker of saccade (Chen-Harris, Joiner, Ethier, Zee, & Shadmehr, 2008) planning. Similarly to saccade amplitude, we also found peak velocity to be biased towards prior target velocity. Peak velocity tended to be higher, compared to control trials, during catch trials with higher prior target velocity, while it tended to be lower when prior target velocity was lower (main effect of trial type; Noisy Target, catch15: F(1,12)=24.05, p=0.0004, ges=0.26, catch20: F(1,12)=12.95, p=0.0037, ges=0.10; Standard target, catch15: F(1,12)=5.36, p=0.04, ges=0.05, catch20: F(1,12)=14.5, p=0.0025, ges=0.17).

### Saccade amplitude is also biased by visual information about the target velocity

As a sanity check for our analysis of saccades, we compared saccades made within conditions having the same prior information and different visual information about the target velocity. As expected, saccade amplitude was also influenced by visual information (Fig. 4), meaning that catch-up saccades accounted for the velocity change. Catch trials with a higher target velocity than control trials (green traces) had larger saccade amplitude, and the opposite pattern was seen for catch trials having a lower target velocity (red traces). Once more, this effect was present for all velocities, and in both experiments (main effect of trial type, Noisy Target: F(2,24)=69.39, p<0.0001 Huynh-Feldt corrected, ges=0.4; standard Target: F(2,24)=211.96, p<0.0001 Huynh-Feldt corrected, ges=0.7). We found no influence of the control (previous) target velocities (15°/s and 20°/s) on the magnitude of the effect (cf. dashed lines vs full lines in Fig. 4).

**Figure 4:**
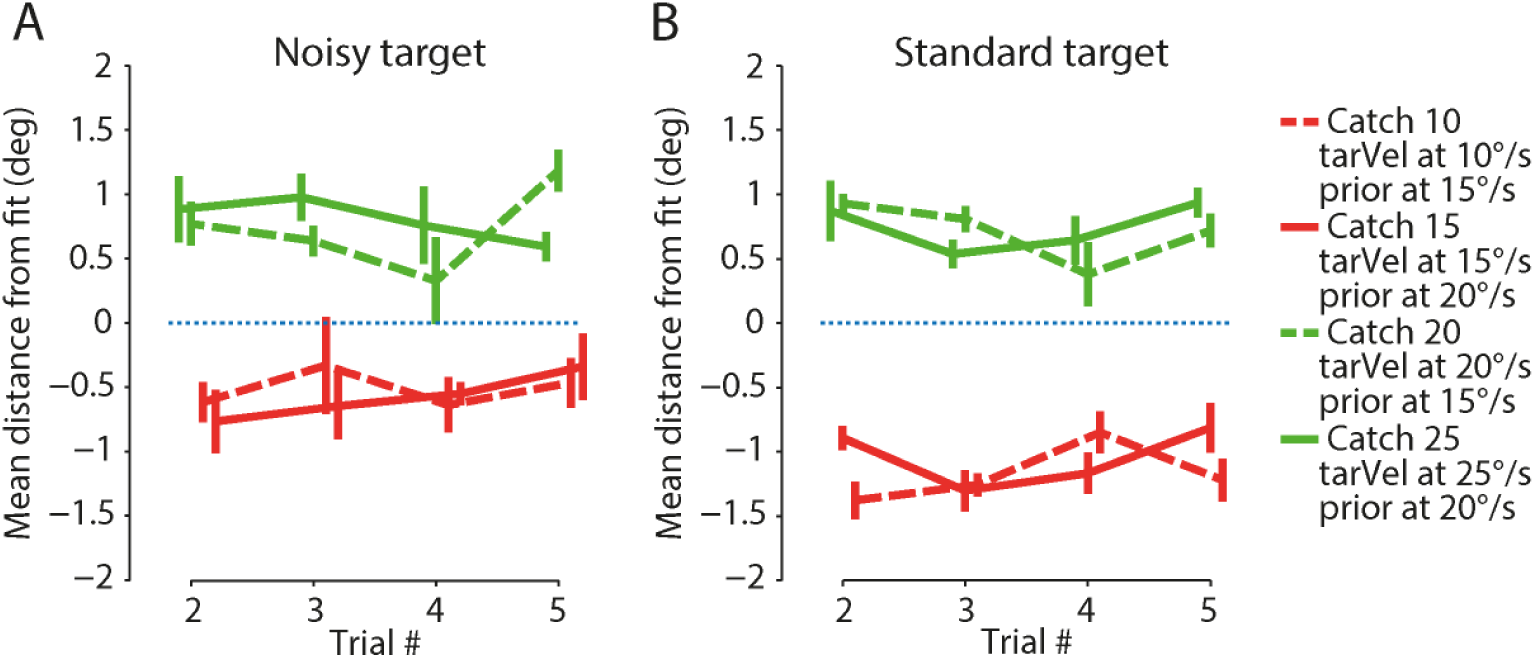
A. Effect of visual information on saccades. Averages of participant’s mean residuals (vertical distance from fit on control data) of the amplitude of saccades made during catch and control trials, with respect to their trial number. Red/green traces show residuals of saccades made during catch trials (the control residuals have been subtracted from all data). The X-axis indicates the trial number and the error bars the standard error of the mean. **B.** Same as panel A, but for the second experiment with a standard target.

### Prior information induces stronger biases when the reliability of the visual information is lower

By comparing the normalized effect of prior information across the two types of target (Fig. 5), we found that the influence of the prior was significantly stronger for the noisy target than for the standard target. As such, the magnitude of the effect on eye velocity was significantly different between the two targets (main effect of target type: F(1,24)=4.43, p=0.046, ges=0.08). Furthermore, we observed significant differences of the modulation of saccade amplitudes by prior information (main effect of target type: F(1,24)=7.17, p=0.0132, ges=0.08). In other words, when confronted to a noisy, less reliable target, participants gave more weight to prior information than when confronted to the standard target.

**Figure 5:**
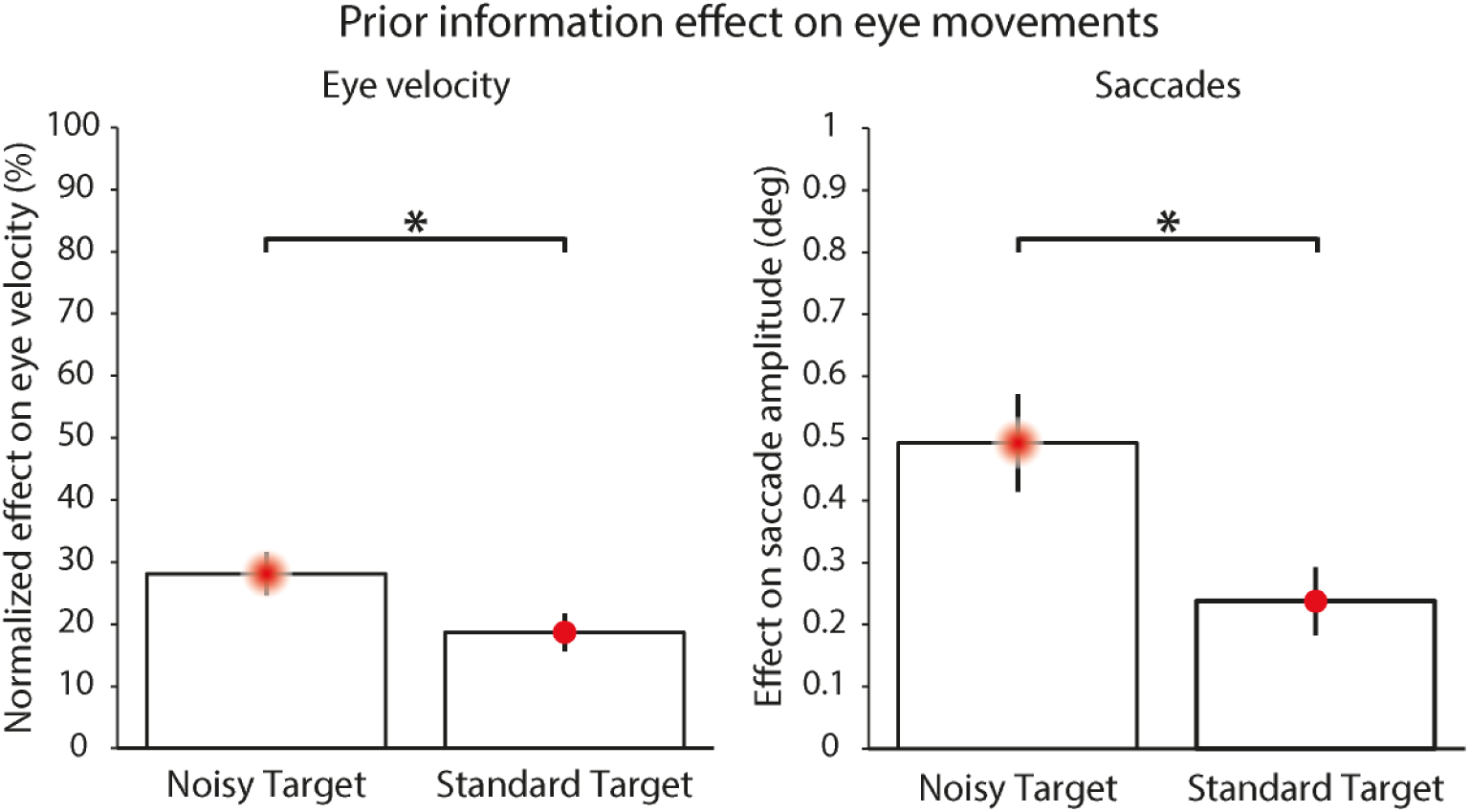
Magnitude of the effect of prior information. Boxes represent averages of participant’s means across both comparable velocities. Effect on eye velocity (left panel, % is relative to the target velocity change: 5°/s) and on saccade amplitudes (right panel). Significativity (* indicates p<0.05) refers to a mixed design ANOVA with within factors ‘target velocity’ and ‘trial#’ and between factor ‘type of target’. Errorbars indicate standard errors of the mean.

### Visual information impact is stronger when the reliability of the visual information is higher

As a complement to the previous analysis, comparing the two types of target (Fig. 6), we found that the standard target elicited a stronger effect of the visual information on both smooth pursuit and saccadic eye movements. Indeed, the effect on eye velocity was stronger (main effect of target type; F(1,24)=20.77, p=0.0001, ges=0.26), and saccade amplitude was more strongly modulated (main effect of target type; F(1,24)=8.64, p=0.007, ges=0.1) when the target was standard rather than noisy.

**Figure 6:**
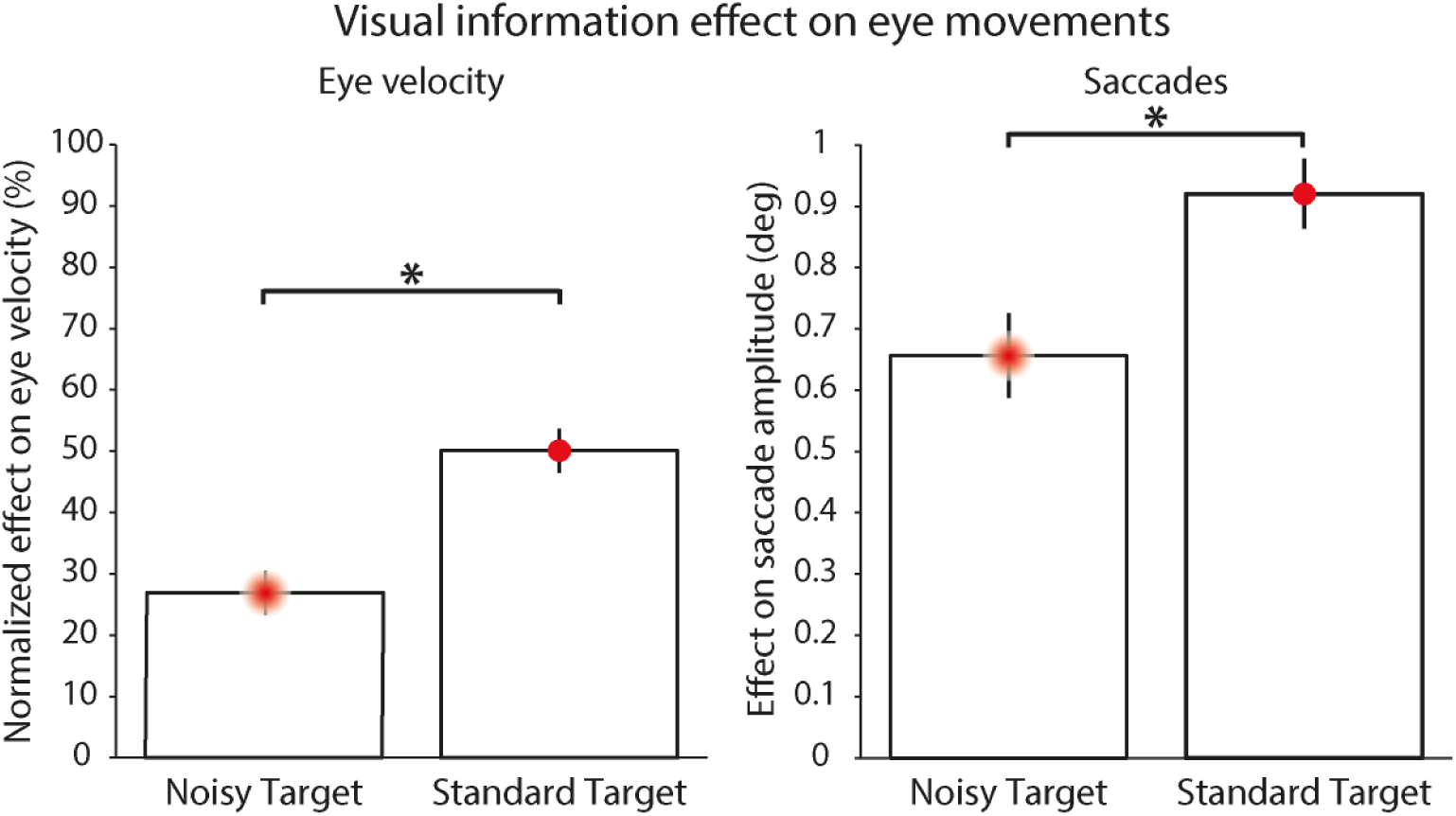
Magnitude of the effect of Visual information on the eye velocity (left panel, % is relative to the target velocity change: 5°/s), and on saccade amplitude (right panel). Boxes represent averages of participant’s means across all comparable velocities. Significativity (* indicates p<0.05) refers to a mixed design ANOVA with within factors ‘control target velocity’ and ‘trial#’, and between factor ‘type of target’. Errorbars indicate standard errors of the mean.

This behavior, opposite to that of prior information, is coherent with the hypothesis that the reliability of visual information is higher for the standard target than for the noisy target, which would give more weight to visual information when pursuing the standard target.

### Higher prior reliability induces stronger effect of the prior, but doesn’t seem to affect visual effect

Postulating that the reliability of the prior information increases with the number of repetitions (Orban de Xivry, Coppe et al., 2013), we analyzed how the effect of prior information (measure of normalized gains, see methods) evolves with trial number. To do so, we computed a simple linear regression to predict the effect magnitude based on the trial number.

Across target types, we found that the magnitude of the effect of the prior on the eye velocity gain significantly increased with the trial number (significant regression slope of 4.9%/trial: F(1,206)=11.39, p=0.0009, R^2^=0.0478), supporting the hypothesis that the weighting of prior information is dynamically updated to reflect its reliability (Fig. 7A). We also computed the linear regression separately for each target type, and observed that for both targets there was a significant increase of the effect with the trial number (noisy target: slope of 4.7%/trial, F(1,102)=5.1, p=0.026, R^2^=0.04; standard target: slope of 5.2%/trial, F(1,102)=6.76, p=0.011, R^2^=0.053).

**Figure 7:**
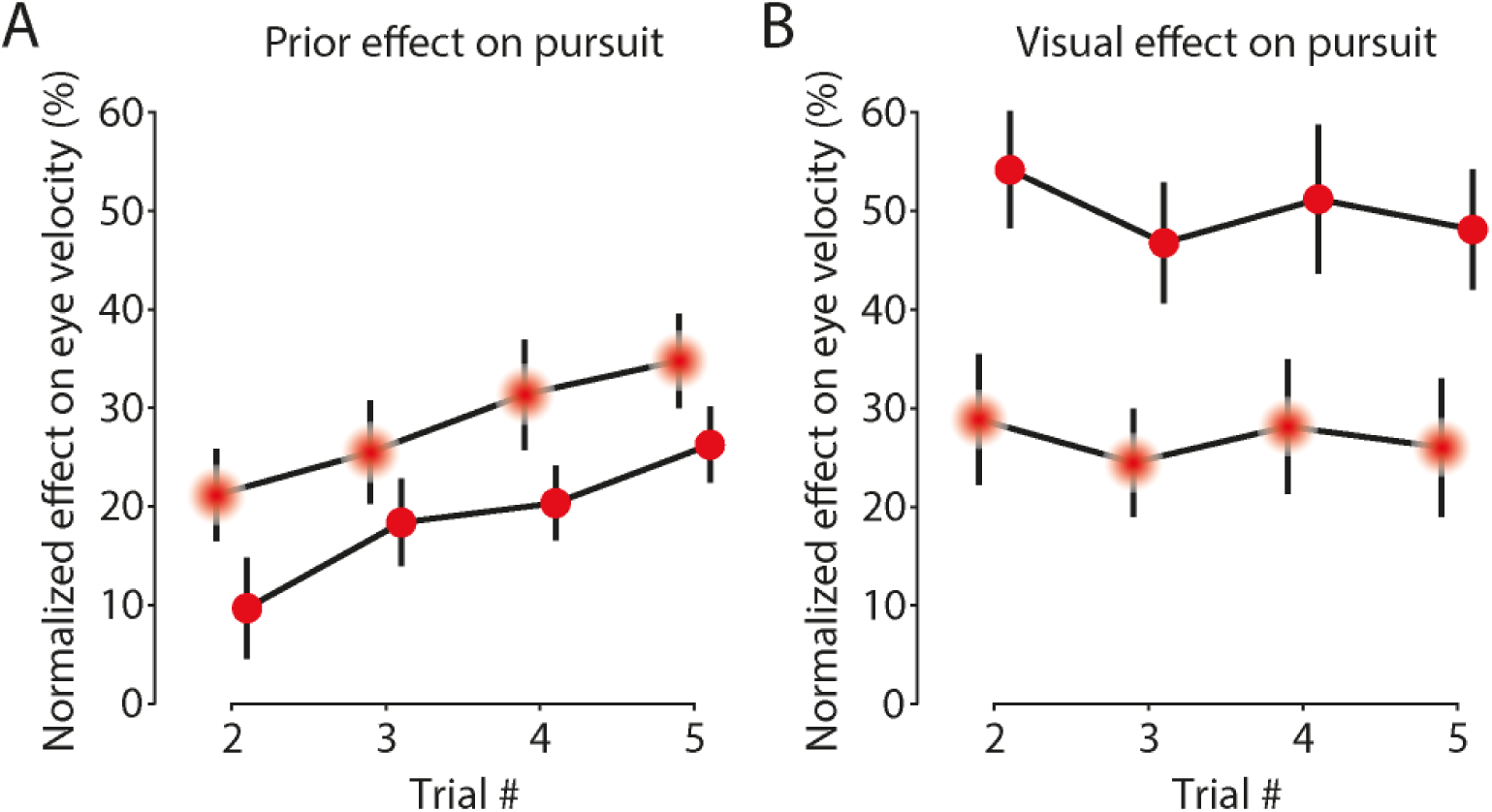
Effect of trial number. **A.** Influence of trial number on the magnitude of the effect of prior information on eye velocity (averages of participant’s means). **B.** Influence of trial number on the magnitude of the effect of visual information on eye velocity (averages of participant’s means). Percentages are relative to the target velocity change (% of 5°/s). Errorbars indicate the standard error of the mean.

In contrast, we found no indication (p>0.39) of an effect of the number of repetitions on the impact of visual information on eye velocity (Fig. 7B).

Note that we could not perform the analysis of interaction of the number of repetitions on saccades because, since we had to match saccade latency distributions to compare saccades across targets (experiments), the number of comparable saccades was reduced by almost 50%. Consequently, the number of comparable saccades per (catch) trial number was too low for such an analysis.

## Discussion

To investigate the dynamic reliability-based-weighting of visual information and memory of target motion during eye movements, we asked participants to visually track either a noisy (Gaussian) target or a standard target for a variable number of identical trials. Then, we tested how the nervous system weighted new sensory information and prior expectations by presenting a trial with a different target velocity.

Our results show that previous trials had a greater effect on eye velocity and catch-up saccades amplitude when the target was a Gaussian blob, in a manner consistent with a reliability-weighted integration of visual information with short-term memory. It was also observed that the effects of previous trials on eye movements appeared as early as after one trial and increased with the number of trials, hinting at a short-term memory of perceived target motion that is dynamically built and updated.

### Eye movements use a reliability-based representation of target motion

It has been known for a long time that extra-retinal and sensory signals can both drive smooth pursuit (Barnes & Asselman, 1991; Dodge et al., 1930; Kowler & Steinman, 1979); however, how these two signals are integrated within the pursuit system has remained elusive. The first models including predictive smooth pursuit had separate pathways which were activated by switches (Barnes, 2008). Such models often struggled to explain the transition between predictive and reactive pursuit and why predictive signals are observed despite the randomization of target features (Kao & Morrow, 1994; Kowler & McKee, 1987). More recently, Orban de Xivry, Coppe et al. (2013) as well as Bogadhi, Montagnini and Masson (2013) both proposed models of smooth pursuit that were able to combine sensory and extraretinal signals without the need for a switch between pathways. Those two models included 2 separate recursive Bayesian/Kalman filters (one for visual inputs, one for extraretinal inputs) and could reproduce many aspects of smooth pursuit behavior in different contexts of visual information. They differed in a few aspects, including the type of memory of the target and assumptions about the visual reliability of the target. Here, we mainly refer to the model of Orban de Xivry, Coppe et al. because we can compare its predictions of the repeated tracking of a noisy target to our experiment. However, given a few modifications, we believe that similar predictions could be made by the model of Bogadhi, Montagnini and Masson.

Compared to the predictions of the model of Orban de Xivry, Coppe et al., we found that our results supported them. Indeed, the modulation of the effect of previous trials on eye velocity by both the reliability of visual information (noisy or standard) and the reliability of the memory (number of repetitions) are strong indicators of a reliability-based integration of visual information with some memory of target motion.

Importantly, our results go further and show that similar processes are at work for catch-up saccades, which fits well with the idea that both saccades and smooth pursuit are influenced by common inputs (Krauzlis, 2004, 2005; Orban de Xivry & Lefèvre, 2007). While these common inputs were either sensory signals from position and motion pathways (Blohm, Missal, & Lefèvre, 2005; S. de Brouwer, Yüksel, Blohm, Missal, & Lefèvre, 2002; Krauzlis, 2004) or from the forward model (Ego, Yüksel, Orban de Xivry, & Lefèvre, 2016; Orban de Xivry, Bennett, Lefèvre, & Barnes, 2006), we show here that the internal representation of target motion, stored in short-term memory, is also shared by the two systems. Hence, catch-up saccade amplitude was affected both by the target velocity of previous trials and by the velocity of the ongoing target. Moreover, the magnitude of the effect of previous trials was greater when the visual target was the (less reliable) Gaussian blob, which is fully compatible with reliability-based integration.

Therefore, we show in the present study that the whole oculomotor system has access to a single short term memory of target motion, and that reliability-based integration captures well the process by which this memory is combined with sensory signals.

### Nature of the representation: short-term memory or prior

Here, we made the assumption that the extra-retinal signals consist of an internal representation of target motion (Orban de Xivry, Missal, & Lefèvre, 2008) and not a prior on target velocity (Bennett & Barnes, 2004; Bogadhi et al., 2013; Yang et al., 2012). These two possibilities differ in the sense that an internal representation of target motion is a memory of the target motion paired to a measure of uncertainty while the prior is a Gaussian distribution with a mean and standard deviation (see Tassinari et al., 2006 for a study on the ability of humans to do reliability-based integration). In addition to previous evidence in favor of an internal representation of target motion (Orban de Xivry et al., 2008), the rapid change in the weight of the extra-retinal signals with trial number (Fig. 7) is at odds with a prior on target velocity, which is commonly slowly built and gradually updated. An internal representation of target motion can also drive anticipatory pursuit (Orban de Xivry, Coppe et al., 2013) and, through continuous reliability-based integration, would fit well the results of Heinen, Badler and Ting (2005) on the effect of previous trials on anticipatory pursuit, and the results of Maus, Potapchuk, Watamaniuk and Heinen (2015) who showed that anticipatory eye velocity was best correlated with the target velocity of the 2 previous trials.

Indeed, we observed that the weight of the extra-retinal signals was updated on a trial-by-trial basis. Such a rapid update in reliability can be modelled through Kalman filtering (Orban de Xivry, Coppe et al., 2013) in the case of an internal representation of target motion. In contrast, achieving the same with a prior on target velocity would require one trial to very rapidly affect its Gaussian distribution, while many trials are usually required to obtain a prior distribution (Yang et al., 2012: several days of training in monkeys; Kording and Wolpert, 2004: 1000 trials in humans). However, there appears to be a trend towards describing shorter timescales for the building of certain sensorimotor ‘priors’, for example of target position: in 2008, Izawa and Shadmehr suggested continuous prior integration with sensory information; in 2011, Verstynen and Sabes reported on ‘fast adapting priors’ built within 10 trials; in 2012, Rao, De Angelis and Snyder suggested rapidly-varying priors downstream of the sensory representation. Priors of target velocity and timing were also suggested to be continuously integrated with other sensorimotor signals by authors such as Heinen, Badler and Ting (2005). Such priors would still allow for other priors built on longer timescales, like priors towards zero target velocity, to be taken into account (Yang et al., 2012) but also explain trial-to-trial effects such as those observed in this experiment.

### Reliability-based integration of a short-term memory with new information

In recent years, reliability-based integration of sensory inputs with other signals has provided an elegant framework to explain how the brain can process noisy and changing visual information to produce appropriate movements. Reliability-based integration has already been observed for inputs from different sensory systems (Ernst & Banks, 2002; Landy et al., 2011), for visual information and ongoing motor commands (Kording & Wolpert, 2004), for visual information and a low-velocity prior (Jogan & Stocker, 2015; Stocker & Simoncelli, 2006) and for sensory predictions and sensory feedback (Vaziri, Diedrichsen, & Shadmehr, 2006) but, to the best of our knowledge, it has until now not been shown to occur for ongoing sensory information and a working memory.

Outside of the oculomotor context, several authors have studied the dynamics of memory and learning and found processes that relate to reliability-based integration of working memory with new information. They have suggested different models for which a memory is updated on the basis of the outcome of a previous action. The core principle is as follows: maintain states/beliefs about the environment (in our case the target motion), and update the existing beliefs on the basis of prior information and recent outcomes for the next trials. In addition, these models particularly focus on whether a new state (a memory) should be formed or the previous one updated, depending on whether a fundamental change in the system is thought to have occurred or not. The states are therefore updated or created depending on the probability of a fundamental change and on the reliability of the signals, namely memory or new sensory information, for example by means of an approximately Bayesian delta-rule model (Nassar et al., 2010; Wilson, Nassar, & Gold, 2013), or a similar optimal filtering model (Gershman et al., 2014). Such belief-updating models have also been applied to the motor adaptation domain (Kording, Tenenbaum, & Shadmehr, 2007; Wei & Kording, 2010).

In all these studies, memory updating is influenced by the reliability of the new incoming information but takes place during the inter-trial interval. In contrast, the present study demonstrates that the integration of the memory content and the sensory information takes place during the movement and directly influences it. Such reliability-based integration of two different signals during movement was reported in a study in which one signal came from the internal model of the arm and the other signal was sensory in nature (Kording & Wolpert, 2004) but, to our knowledge, it is the first time that reliability-based integration during movement is reported for sensory and working memory signals.

### Integration of short-term memory and visual signals in other contexts

Several studies investigated the influence of the memory of a target position on reaching arm movements. Brouwer and Knill (2007, 2009) showed that reaching movements towards a visual target were biased towards its last known position and that the effect was stronger if the contrast of the target was low. Verstynen and Sabes (2011) also showed the presence of a bias towards previous positions of the target during reaching movements despite the fully predicable nature of the next positions. In addition to the position of the target, the spatial structure of the environment itself has also been shown to be memorized during visuomotor tasks (Aivar, Hayhoe, Chizk, & Mruczek, 2005; Hayhoe, Shrivastava, Mruczek, & Pelz, 2003).

Those studies clearly highlight that the reliability of a visual target, and previous information about it, can affect hand movements in a way that is similar to their effect on eye movements (Issen & Knill, 2012). However, these studies mainly focused on the integration of a memory of target position with the visually presented one. Such memories are limited to a position signal and a spatial representation of the scene and do not contain the evolution of the signal over time. Given that we know that the oculomotor system does not restrict this memory to a measure of target velocity but also includes its time course (Bennett, Orban de Xivry, Lefèvre, & Barnes, 2010; Orban de Xivry et al., 2008), we believe that the content of the working memory used for reliability-based integration during movement is much more complex than the ones previously described (Song & Nakayama, 2009).

Finally, while the integration of the position signals from visual information and memory occurs mostly before movement onset, we observe here a continuous integration of working memory and sensory information during eye movements. Given the similarities between the oculomotor system and other sensorimotor systems (Lisberger, 2015; Lynch & Tian, 2006), we may expect to find a similar process of continuous short-term memory updating in those systems.

## Conclusion

In this study, we report experimental evidence in the context of oculomotor behaviors that short-term memory can be quickly built, constantly updated and continuously integrated in a reliability-based manner with incoming visual information. We believe that this constitutes a general principle of dynamical updating of working memory, one that is consistent with two recent studies (Gershman et al., 2014; Nassar et al., 2010), and that is likely to be present in other sensorimotor systems (Lisberger, 2015).

## Methods

### Participants

Because of the absence of previous literature on the topic, a power analysis could not be used beforehand to determine the number of participants. We therefore decided to refer to what is typically done in eye movement research and targeted a pool of more than 10 participants per experiment. Twenty participants between 18 and 30 years old were recruited to participate in our experiments. Thirteen participants (4 female) participated in the first experiment and thirteen (6 from the first) in the second experiment (5 female).

Participants had normal or corrected to normal vision. After being given a full description of the experiment, informed consent was given by the participants. The procedures were approved by the Université catholique de Louvain Ethics Committee and were in accordance with the Declaration of Helsinki.

### Protocol

Participants were seated in a dark room, and looked at a 197x150 cm screen at 151 cm in front of them, spanning ±40° of their visual field. Head movements were restrained with chin and forehead rests. The stimuli were projected onto the screen with a cine8 Barco projector (Barco Inc., Kortrijk, Belgium) at a refresh rate of 100 Hz and the eye movements were recorded at 1000Hz using an Eyelink 1000 (SR Research, Ottawa, Ontario, Canada). The display of visual stimuli was handled by an in-house toolbox, while interactions with the Eyelink^®^ were handled by the Psychtoolbox (Kleiner et al., 2007). Calibrations trials were first performed at the start of the experiments, then every 30 trials (±2min). Breaks were allowed before every calibration (every 2min), and the total duration of an experiment was around 30 minutes.

In the design of the protocol, we wanted to make sure that behaviors could only be related to the current block, i.e. that there was no transfer of information between blocks. As such, block duration and features were randomized, direction changed after each block and each of them started with a passive trial meant to wash out previous block-related memories.

Two types of stimuli were used: a red fixation target (uniform disk, diameter 0.8°), and a red pursuit target. The protocol was identical for both experiments except for the pursuit target. In experiment #1, the pursuit target was a 2D Gaussian spot (σ=1.27°; hereafter the *noisy* target, cf. Fig. 8 - supplement 1). In experiment #2, the pursuit target was a uniform disk (0.8° of diameter; hereafter the *standard* target, cf. Fig. 8 - supplement 2). The overall luminance of the stimuli was the same for all stimuli, however, given the difference in the distribution (the pixel at the center of the noisy target has 5% of the luminance of any pixel of the standard target), the noisy target was harder to perceive.

For both experiments, there were two main types of trial, *passive* and *active*, and two sub-types (of *active* trials): *control* and *catch*. All trials started with the display of the fixation target at the center of the screen. After 500ms, the pursuit target appeared at the center and immediately moved in one out of 6 possible directions (-20°, 0°, 20°, 160°, 180° or 200°) at a constant velocity (15°/s or 20°/s for control trials) during 650ms. In passive trials, the fixation target remained on for the whole trial, and participants were instructed to keep looking at the fixation target while inhibiting movements towards the pursuit target. In active trials, the fixation target disappeared at the onset of the pursuit target and participants were instructed to follow the pursuit target with their eyes (see Fig. 8).

The trials were presented in blocks: each block consisted of one passive trial followed by 1 to 5 active trials with 850ms between trials. To warn participants of a transition from active to passive trial, i.e. the start of a new block, an auditory cue (440Hz, 80ms) was given 200ms before the appearance of the fixation target. Target direction and velocity remained constant throughout a block, except for the last trial, hereafter named *catch* trial, for which velocity was reduced or increased by 5°/s with respect to previous trials of the same block. Active trials that are not catch trials are hereafter categorized as *control* trials. To summarize, each block started with one *passive* trial and ended with one *catch* trial, with up to 4 *control* trials in-between (Fig. 8B).

There were 120 conditions in total (6 directions x 2 control velocities x 5 block lengths x 2 catch velocities). The average number of trials per condition was 4, yielding 480 trials per participant. The collected data is available from the Dryad Digital Repository (http://dx.doi.org/10.5061/dryad.h0qr3).

### Comparisons between trials

**Figure 8:**
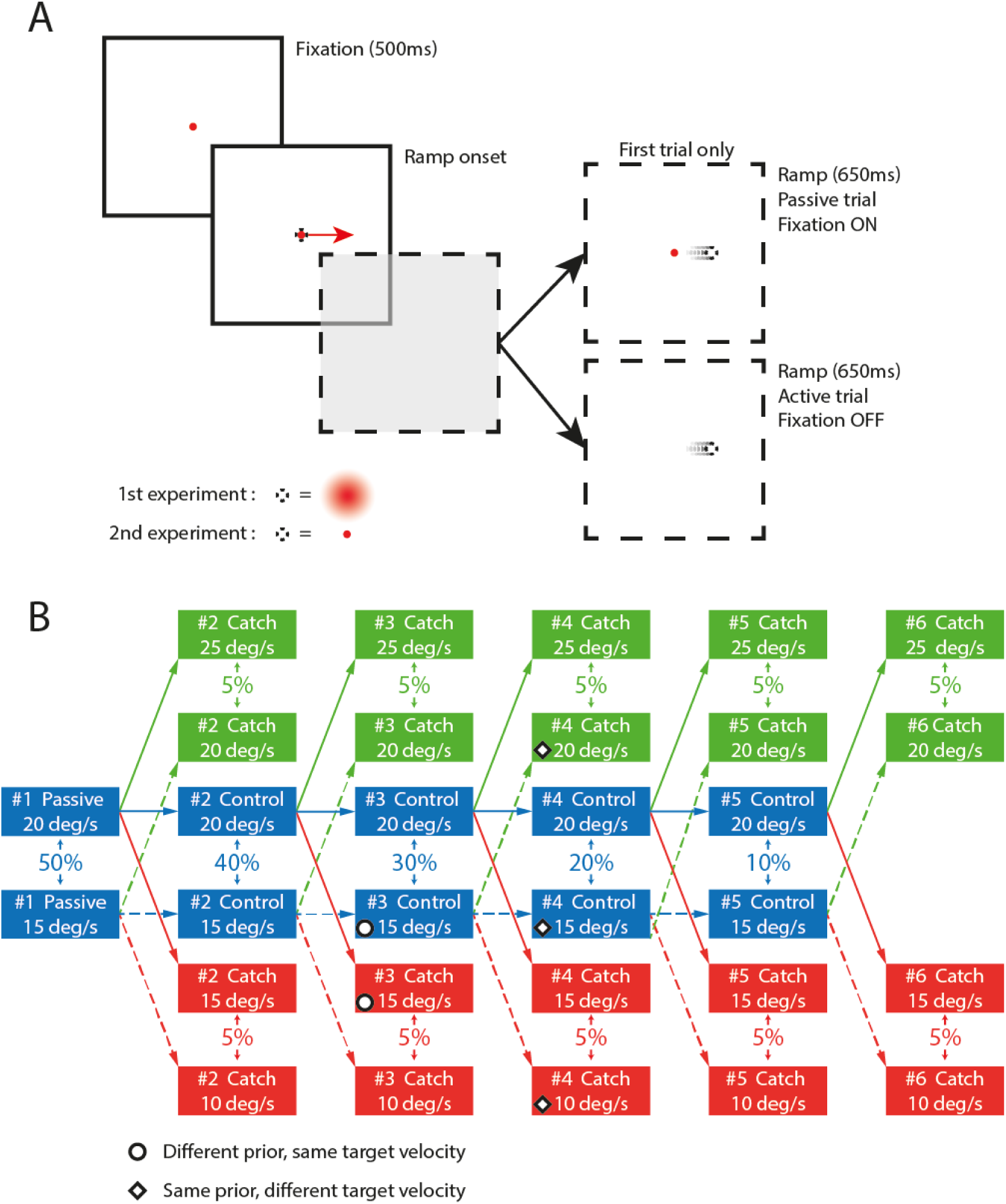
Experimental protocol. **A. Trial example.** Timing of trials (for one of the six possible directions). **B. Template of blocks (regardless of direction).** Each block starts with a passive trial and ends with a catch trial (red/green). Percentages indicate the % of blocks containing trials of the same type (color) and number (#) at one of the two available velocities (ex: 100% of the blocks have one passive trial). Black & white disks give an example of two trials that can be compared to highlight the effect of previous trials. Black & white diamonds give an example of trials comparisons highlighting visual effect. Dashed arrows indicate a 15°/s prior target velocity. Solid arrows indicate a 20°/s prior target velocity. Note that the noisy target shown here has been made more salient to ensure that it remains visible. Its actual appearance is shown in Figure 1 - supplement 1, with supplement 2 showing the standard target as reference.

To observe the influence of previous trials on the oculomotor response, we compared trials that had the same current target velocity, but different past target velocities (different prior trials). For example, we compared *catch* trials having a target velocity of 15°/s (thus preceded by control trials at 20°/s) with *control* trials having a target velocity of 15°/s (thus preceded by control trials at 15°/s). In this situation, visual information (target motion) is the same for both catch and control trials, but prior information is different, effectively highlighting its effect (cf. black & white disks in Fig. 8B).

In a similar way, the impact of visual information was estimated by comparing catch trials to control trials that had the same past target velocity, but different current target velocities. (cf. black & white diamonds in Fig. 8B).

We always made comparisons between trials with the same trial number (in the same column on Fig. 8B).

### Data processing

Data were processed using the Matlab^®^ software (RRID:SCR_001622). Blinks were detected based on missing values in the Eyelink^®^ output (when the pupil cannot be detected) and subsequently removed from the data, including a safety margin before and after the blink, up to the first local minimum in the y-coordinate. Eye position signals were low-pass filtered at 40 Hz. Eye velocity and acceleration were obtained from position signals using a central difference algorithm on a ±10-ms interval. For the analyses, we pooled data across all directions.

Saccade onset and offset were detected using a 500°/s^2^ threshold on the acceleration data. Saccades were thereafter removed from smooth eye velocity data and replaced by linear interpolation.

In order to remove abnormal trials from the data set while limiting visual inspection of the data, we set a few criteria: (1) during the last 100ms of fixation, eye position within 3° of the fixation target, (2) no missing data (blinks) in the first 450ms of pursuit, (3) lower limit on eye displacement of at least 40% of target displacement, (4) no eye velocity over 40°/s during pursuit epochs. Based on these criteria, we set aside less than 3% of the trials. When analyzing eye velocity during pursuit epochs, we included only trials for which steady state pursuit velocity reached 33.33% of target velocity (98% of trials).

### Eye velocity and pursuit gain computation

Eye movement velocity during steady-state pursuit was obtained by fitting a piece-wise linear regression on the eye velocity data (mean least square regression, using the *lsqcurvefit* function of the Matlab^®^ Optimization Toolbox), as follows (Fig. 9):

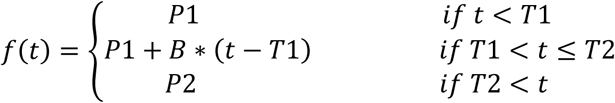

**Figure 9:**
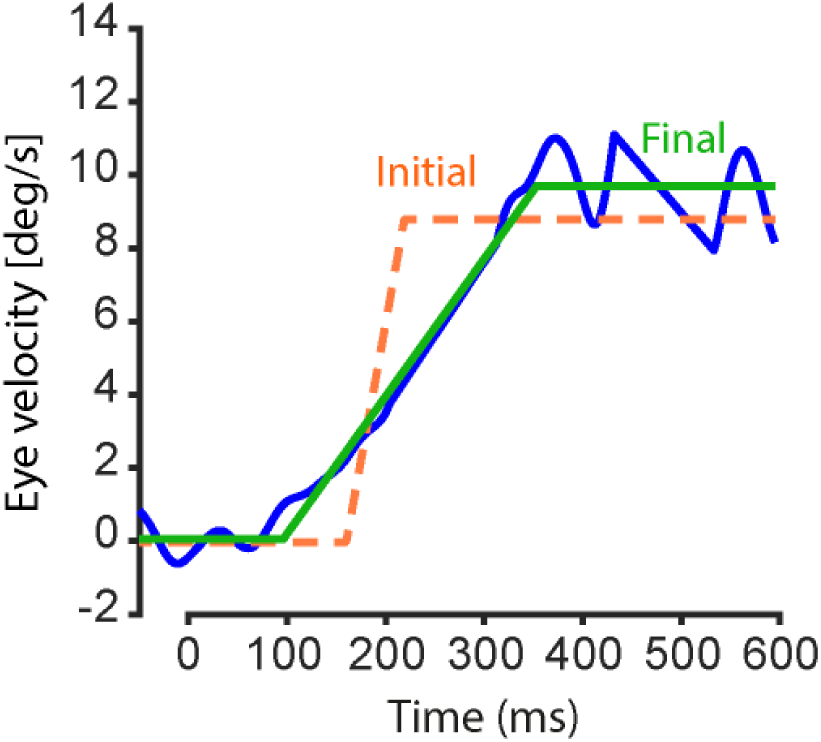
Typical fit and its initial parameters. The orange dashed line shows the initial parameters for the fit. The continuous green line shows the resulting fit of the eye velocity data.

Where *t* is time in seconds, P1 and P2 are velocity plateaus (representing initial and steady-state velocities - in degrees per second), T1 and T2 are the reaction time and the pursuit steady-state onset time (seconds), and B is the initial acceleration, connecting the plateaus between times T1 and T2 (degrees per second squared). P1, P2, T1 and B were the free parameters of this fit. The fitted interval spanned 600ms, from 50ms before target onset up to 600ms after. Initial parameters for P1, P2, T1, and B were determined as follows: P1 was set to the average velocity in the first 100ms of the interval, T1 to the time after which the eye velocity exceeded 20% of the target velocity. The variable T2 was defined as the start of the sub-interval during which eye velocity exceeded 33% of the target velocity for at least 125ms. P2 was then set to the average value of the eye velocity during the interval. Finally, B was determined from the previous parameters, such that it didn’t exceed 150°/s^2^. If any of the conditions couldn’t be met, or if a suitable interval (>33% of target velocity for 125ms) couldn’t be found, the initial parameters were set to default values: T1 to 320ms, T2 to 470ms and P2 to 80% of the target velocity.

Since we wanted the fitting algorithm to measure smooth pursuit eye velocity, we gave less weight to the eye velocity data interpolated during catch-up saccades, setting it to 0.3 (compared to 1 for pursuit data). After applying the fitting algorithm, trials whose steady-state velocity plateau (P2) duration was under 50ms were analyzed again. They went through a second step of fitting, using the same function, but with T2 as a free parameter instead of B. This allowed the use of a different set of initial values, with a steady-state velocity plateau lasting at least 50ms. After the second fitting, any trial whose fitted steady-state plateau (P2) was still less than 50ms long was rejected (±6% of trials), as it meant that the algorithm could not find the steady-state of the smooth pursuit.

We computed the eye velocity gain of a trial by dividing the eye velocity during steady state by the target velocity of a control trial with the same trial number. The control trial used was the one the catch trials were being compared to, and therefore depended on the type of comparison (prior or visual). When studying the influence of prior information, the eye velocity gain was computed with respect to the control trial having the same target velocity as the catch trial (cf. black & white disks in Fig. 8B). When studying the influence of visual information, the eye velocity gain was computed with respect to the control trial having the same prior target velocity as the catch trial (cf. black & white diamonds in Fig. 8B).

When we compared the effect of prior on gains across different velocities, we normalized eye velocity using the following equation (1):

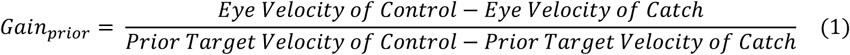

When the comparison across velocities had to be made for the effect of visual information, eye velocity gains were normalized using the following equation (2):

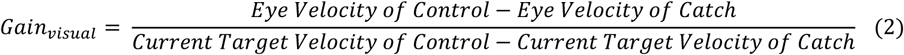

### Saccade metrics

We also studied the amplitude of the first (catch-up) saccade occurring between 100ms and 400ms after target onset (94% of first saccades made after target onset). The amplitude of the catch-up saccades made during control trials was used as a reference to build a linear model of the saccadic behavior of each participant (cf. Fig. 10).

**Figure 10:**
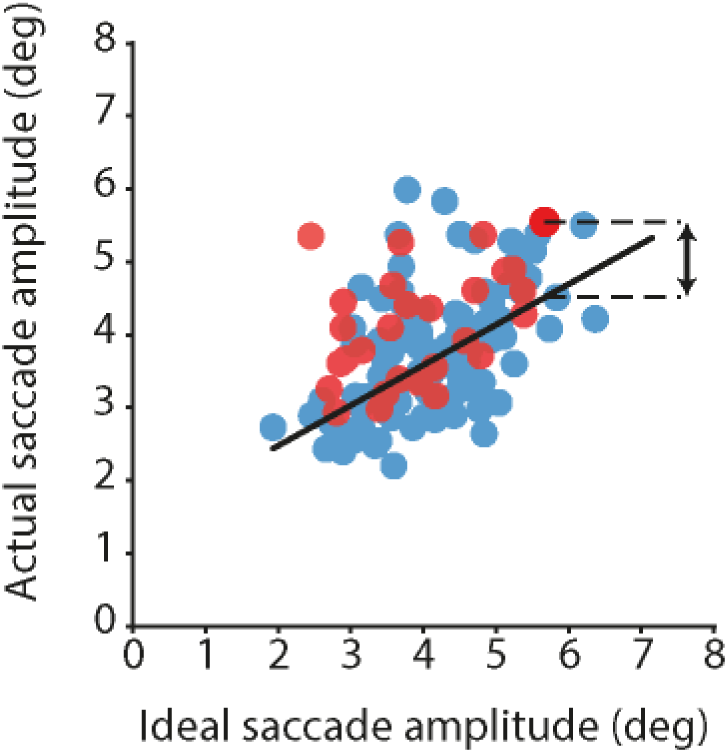
Computation of saccade residuals. Blue dots correspond to saccades made during control trials, red dots to saccades made during catch trials with the same target velocity, but whose prior trials had a higher target velocity. The fit made on the control data is shown by the black line. Dashed lines and arrows show the measure of the residual of one of the catch trials.

For each of the participants, we computed the ideal amplitude of each saccade (difference between eye position at the onset and target position at the offset), and its actual amplitude. Then, the baseline relationship between those two parameters was obtained from control trials by fitting a linear regression (*robustfit* function, Statistics Toolbox, Matlab^®^) on the saccades data. Finally, saccades made during catch trials were compared to this regression line by computing the mean of the residuals (vertical distances between catch saccades data points and the regression line). Given this method, a mean value greater than zero implied that saccades made during catch trials had larger amplitudes than those made during control trials.

To compare saccades made during catch trials across experiments (standard and noisy targets), we had to take into account differences of average saccade latency between the two experiments. Therefore, we used a bootstrap procedure to create N=10000 samples of saccades from the two experiments, such that, for each sample, the latency distributions of the two experiments would be the same. This was done for each control target velocity. For each participant, the average residuals (one per control velocity) for each experiment were then obtained by averaging the average residuals 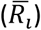 obtained for each of the 10000 bootstrapped samples 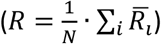. For example, considering the saccades depicted in Fig. 10, the bootstrap procedure might give several subsets of the red dots (catch trials), which will then be compared to the regression model built from all blue dots (control trials) to obtain one average residual per subset.

### Data analysis

The effect of prior information (see previous section) on eye velocity or saccade amplitude was analyzed by comparing trials with the same target velocity but different priors of target velocity. For each *control* target velocity there was only one corresponding *catch* target velocity condition (catch at 20°/s-5°/s for a control at 15°/s, 15°/s+5°/s for a control at 20°/s). For each of those 2 conditions separately, we compared eye movement measures using a repeated measures analysis of variance (rANOVA), using the trial number (4 levels) and the type of trial (control or catch) as within-subjects factors.

The effect of visual information on eye velocity or saccades was analyzed by comparing trials with the same target velocity during the previous trials (15 or 20°/s) but with different target velocities in the current trial (three levels: -5, 0, +5°/s with respect to target velocity during the previous trials, which correspond to 3 types of trial: catch -5°/s, control and catch +5°/s). Eye movement measures were compared using repeated measures ANOVA with the following within-subject factors: target velocity in previous trials (2 levels), type of trial (3 levels) and trial number (4 levels).

When comparing data across the two types of target to examine differences in the magnitudes of the effects, we used a mixed-design analysis of variance with the type of target as between-subject factor. Comparisons on the effect of the prior were made for the two velocity conditions (after normalization with control data), and included the trial number as within-subject factor. The comparison was also made with the velocity condition as an additional within subject factor. Comparisons on the effect of visual information were made with target velocity in previous trials, type of trial and trial number as within-subject factors.

We analyzed the influence of the number of trials on the magnitude of the effects using linear regressions. ANOVAs and linear regressions were performed with the R software (R Core team 2016, RRID:SCR_001905; ez package 2015). Sphericity assumptions were verified through Mauchly’s Test for Sphericity. If sphericity assumptions were violated, we only reported results that were significant after Huynh-Feldt sphericity correction. When appropriate, we also reported the Generalized Eta-Squared (ges) as a measure of effect size (Bakeman, 2005).

## Figure supplements

**Figure 8 – supplement 1:**
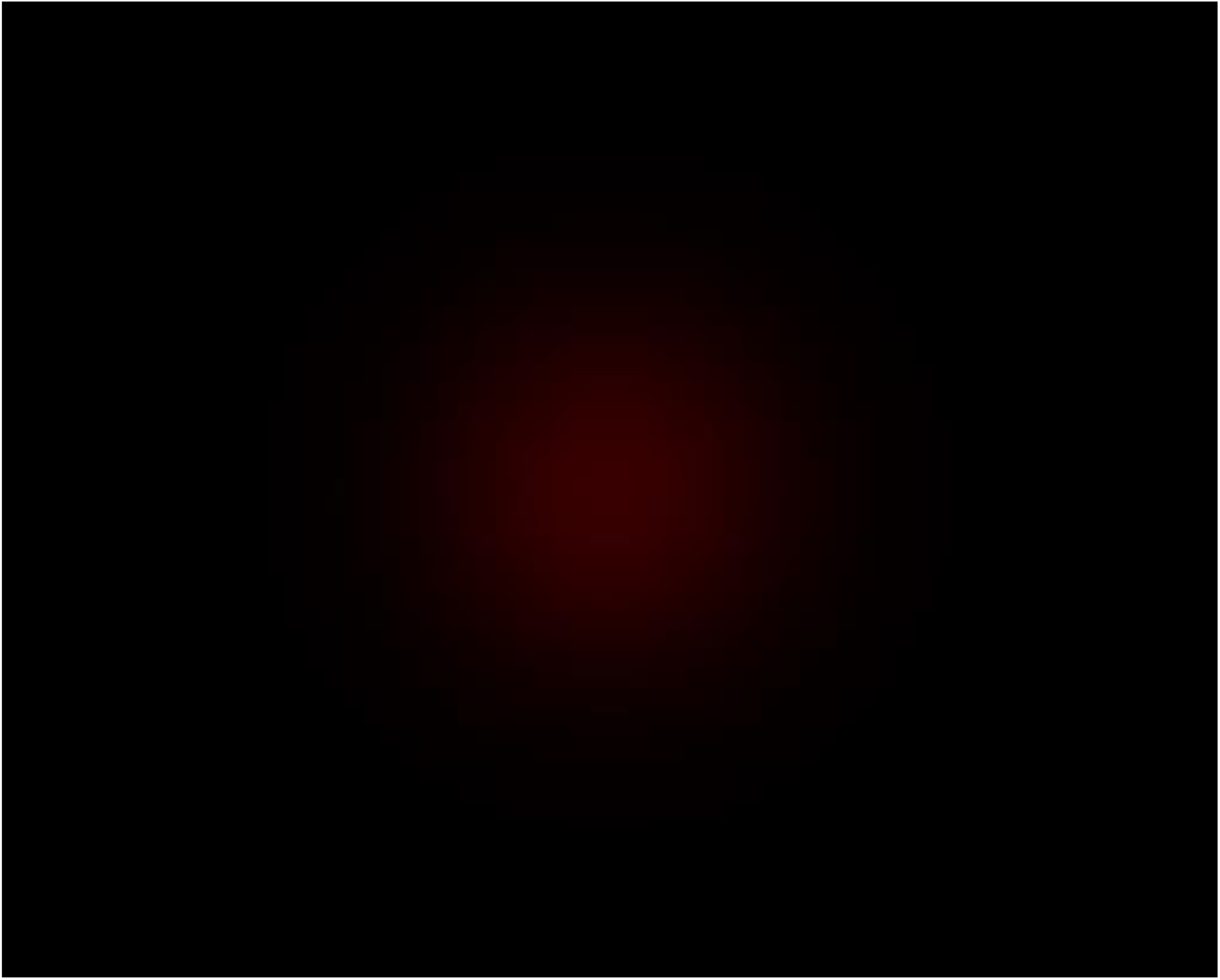
Noisy, 2D Gaussian target on a dark erimental conditions. Note that to obtain a correct representation of its luminosity distribution, it is necessary to correct for the non-linearity in RGB color scaling of the display. If the picture is properly scaled so that its height is 12cm, seeing the target at 50cm should be equivalent to the perspective of the participants.

**Figure 8 – Supplement 2:**
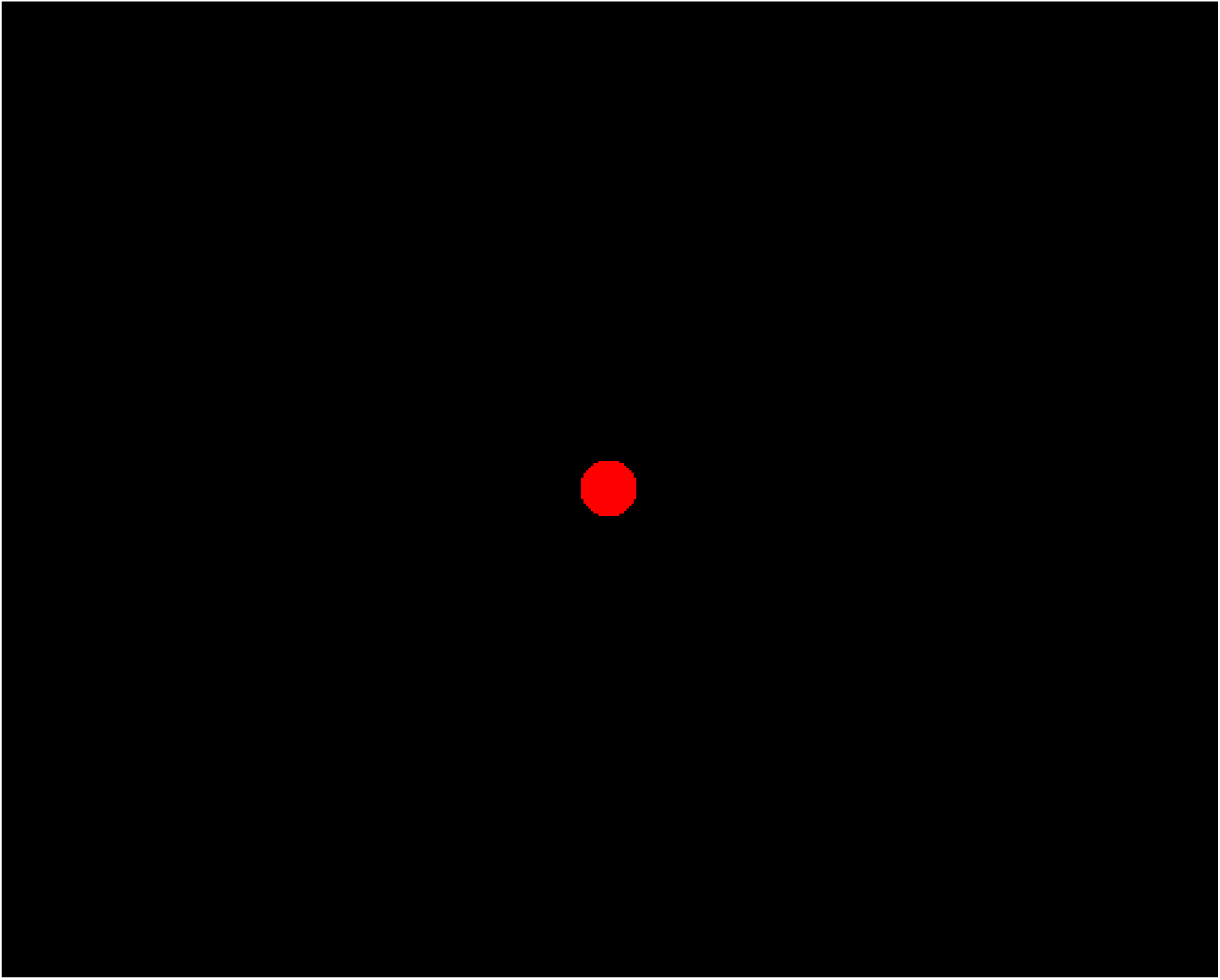
Standard target (0.8° of diameter). If the picture is properly scaled so that its height is 12cm, seeing the target at 50cm should be equivalent to the perspective of the participants.

